# A neuronal thermostat controls membrane fluidity in *C. elegans*

**DOI:** 10.1101/2019.12.20.882514

**Authors:** L Chauve, S Murdoch, F. Masoudzadeh, F. Hodge, A. Lopez-Clavijo, H. Okkenhaug, G. West, A. Segonds-Pichon, S. Wingett, M. Wakelam, K. Kienberger, K. Kleigrewe, O Casanueva

## Abstract

An organisms’ ability to adapt to heat can be key to its survival. Cells adapt to temperature shifts by adjusting lipid desaturation levels and the fluidity of membranes in a process that is thought to be controlled cell autonomously. We have discovered that subtle, step-wise increments in ambient temperature can lead to the conserved heat shock response being activated in head neurons of *C. elegans*. This response is exactly opposite to the expression of the lipid desaturase FAT-7 in the worm’s gut. We find that the over-expression of the master regulator of this response, Hsf-1, in head neurons, causes extensive fat remodeling to occur across tissues. These changes include a decrease in FAT-7 expression and a shift in the levels of unsaturated fatty acids in the plasma membrane. These shifts are in line with membrane fluidity requirements to survive in warmer temperatures. We have identified that the cGMP receptor, TAX-2/TAX-4, as well as TGF-β/BMP signaling, as key players in the transmission of neuronal stress to peripheral tissues. This is the first study to suggest that a thermostat-based mechanism can centrally coordinate membrane fluidity in response to warm temperatures across tissues in multicellular animals.

## Introduction

The ability of an organism to adapt to environmental change can be key to its survival. The model organism *C. elegans* survives and reproduces optimally over an environmental temperature range of 12°C to 25°C (Sulston JE 1988). Temperatures beyond this range cause *C. elegans* severe physiological stress(van Oosten-Hawle and Morimoto 2014). When exposed to heat stress, *C. elegans* activates a highly conserved stress response, called the Heat Shock Response (HSR), during which the Heat Shock Factor 1 (HSF-1) transcription factor rapidly induces the expression of heat shock proteins (HSPs) (Lindquist 1986; Åkerfelt, Morimoto, and Sistonen 2010), which then clear and refold heat-damaged proteins (Morimoto 1997).

In addition to such cell autonomous responses, centrally controlled strategies also help to integrate external and internal cues across tissues to regain homeostasis. Rats, like other mammals, are endotherms and can control their body temperature. They have Transient receptor potential (TRP) channels in their skin and abdomen, which detect ambient temperature. This information is integrated in the hypothalamus, which serves as a thermostat (Madden and Morrison 2019). The efferent output from the hypothalamus provides negative feedback by controlling heat dissipation when it is warm, and heat conservation and thermogenesis when it is cold (Madden and Morrison 2019). Invertebrates, such as *drosophila* and *C. elegans*, are ectotherms and do not internally regulate temperature. However, they possess well-described thermostat-mediated escape responses that enable it to avoid noxious stimuli, for example, the heat-dependent activation of the bilateral thermosensory AFD neurons, which coordinate a thermotaxis response (Kimura et al. 2004; Hedgecock and Russell, 1975)

Although thermostat-based behaviours help ectotherms to escape noxious stimuli like heat, the longer-term survival of ectotherms when environmental temperatures change depends on their ability to remodel lipids within the plasma membrane, which is highly sensitive to external temperatures. Homeoviscous adaptation (HVA) is a mechanism that regulates the viscosity and permeability of membranes to ensure the robustness of biochemical reactions(Sinensky 1974; Cossins and Prosser 1978). In homeoviscous cold adaptation, membrane bilayers undergo a reversible change of state from a non-fluid (ordered) to a fluid (disordered) structure whereby the membrane’s phospholipids (PLs) fatty acyl (FA) chains become increasingly unsaturated (Mendoza 2014; Ernst, Ejsing, and Antonny 2016). In *C. elegans*, three Δ9-acyl desaturase enzymes, FAT-5, FAT-6 and FAT-7, *de novo* synthesize FAs, and in particular monounsaturated FAs (MUFAs)(Watts and Ristow 2017). These FAT enzymes are orthologs of human stearoyl-coA desaturases (SCDs) (Brock et al, 2006a). Of these three *fat* genes, *fat-7* is upregulated at 16°C and is essential for maintaining the fluidity of membranes at cold temperatures (Murray et al. 2007; Savory et al, 2011a; Taylor et al., n.d.; Shi et al. 2013). In warm temperatures, the opposite occurs and *fat-6/7* are negatively regulated (Lee et al. 2019a) concomitant with increased FA saturation in PLs (Tanaka et al. 1996; Murray et al, 2007). Indeed, mutants that abnormally upregulate *fat-7* cannot survive at 25°C (Ma et al, 2015).

HVA was first discovered in single-celled organisms in which membrane sensors control membrane fluidity(Ernst et al, 2016). In *C. elegans*, two temperature-controlled cell autonomous sensors adjust FAT-7 expression: the transmembrane cool-sensor PAQR-2/AdipoR2 (Svensson et al. 2011; Svensk et al. 2013) and the heat-induced Acyl-CoA dehydrogenase, ACDH-11(Ma et al. 2015). PAQR2 reportedly also has a cell non-autonomous role in modulating HVA (Svensk et al. 2013) but no direct input from the brain has yet been linked to this adaptive response.

Increasing evidence indicates that across metazoans, neurotransmitters and neurohormonal signals modulate fat metabolism across tissues (Mattila and Hietakangas 2017; Dietrich and Horvath 2012; Cornejo et al. 2016; Srinivasan et al, 2015). In *C. elegans*, many of these pathways converge on the regulation of fat stores in the gut by controlling either the expression of lipases (Noble, et al 2013) or the *Fat-7-*dependent *de novo* synthesis of FAs (Yu et al. 2017; Clark et al. 2018; Horikawa and Sakamoto 2010) . Notably, ligands from the TGF-β/BMP signalling pathway provide a good example of this regulation; these ligands are secreted from neuronal cells and lead to peripheral fat remodeling in fat store cells(Yu et al. 2017). However, this well-described molecular pathway, which is also conserved in mammals (Tan et al. 2012), has not been studied in the context of HVA.

Here, we explore the antagonistic relationship between the Hsf-1-mediated activation of the HSR and the regulation of unsaturated lipids in *C. elegans*. Previous experiments have shown that while HSP levels increase in ambient temperatures of 25°C compared to 15°C, SCD enzymes show reduced expression at this temperature (Savory eta al 2011a; Lee et al. 2019a) . In addition, both the regulators of SCD enzymes-such as MDT-15- and SCD enzymes themselves are known to negatively regulate HSP expression at 15°C^19^(Savory et al 2011a) . However, it is not known whether Hsf-1 dependent cellular responses can modulate SCD expression and MUFA levels. Here, we show that *hsp* transcripts are expressed primarily in the head neurons of *C. elegans* at 20°C and show a clear temperature-sensitive expression pattern. We used the neuronal overexpression of HSF-1 (*nhsf-1)* as a tool to study the consequences of ectopically activating HSR in neurons at 20°C. We find that HSF-1 overexpression, via *nhsf-1*, in addition to its previously described role in controlling peripheral stress responses and longevity (Douglas et al. 2015), remodels lipid metabolism. This function is performed by decreasing the expression of *fat-7* in the intestine, whilst activating the expression of catabolic lysosomal lipases. We identified the cGMP receptors TAX-2/TAX-4 and TGF-β/BMP signaling to be essential for the transmission of neuronal heat stress information to the intestine. We also show that *nhsf-1* can turn on a similar fat remodeling program as that used by nematodes raised at 25°C, resulting in lower oleic acid levels in membrane phospholipids and a general decrease in FA saturation. This is the first study to report that ectotherms might use thermostat-based mechanisms to centrally coordinate complex adaptive responses to warming temperatures, as occurs in endotherms.

## Main text

### Fat desaturase expression negatively correlates with heat shock protein expression at increased temperatures

*Hsp* genes are expressed more highly at 25℃ relative to their expression levels at 20℃ or 16℃ in *C. elegans* (Gomez-Orte et al, 2018; Lee et al, 2009; Lee et al, 2019). Nevertheless, their overall expression levels are low even at 25℃, precluding the traditional *in vivo* visualization of multi-copy *Hsp* transgenic lines (Guisbert et al. 2013). To circumvent this technical issue, we took advantage of cGAL, a temperature-robust GAL4-UAS binary expression system, which is a more robust gene expression system than traditional transcriptional reporters (Wang et al. 2017). As shown in **Figure 1A** and B, When GAL4 is driven by an *hsf-1* dependent promoter (*hsp16.41*) in *C. elegans*, GFP expression is primarily restricted to cells that have axonal projections in the head region. Surprisingly, GFP is only expressed in neuronal cells, and not in any other somatic tissues (**Figure 1B**), indicating that head neurons are exquisitely sensitive to small, step increments in temperature. Single-cell RNA sequencing confirmed that the only tissue with detectable *hsp* levels are head neurons (Taylor et al., n.d.). We observed that the number of GFP-positive neurons increased almost 16-fold from 15℃ to 25℃, when *gfp* was driven by *hsp16.41*(**Figure 1C, Supp Table 1A**). To rule out that this might be due to an indirect effect of temperature on the expression capacity of the GAL4-UAS binary system, we looked at the output of *unc-47*, a promoter that is restricted to GABAergic neurons but that does not have HSF-1 binding sites. As shown in **Supp.** **Figure 1A** **and Table S1A**, and as expected for a temperature-robust cGAL system (Wang et al. 2017), we observed that when *gfp* is driven by *unc-47*, there is no significant temperature-dependent shift in the expression of *gfp*. A positive interaction measured by a two-way ANOVA analysis indicates that the temperature-sensitive behavior of the *hsf-1* responsive promoter is not driven by the effect of temperature on the binary expression system but reflects the activity of *hsp16.41*.

**Figure 1.**
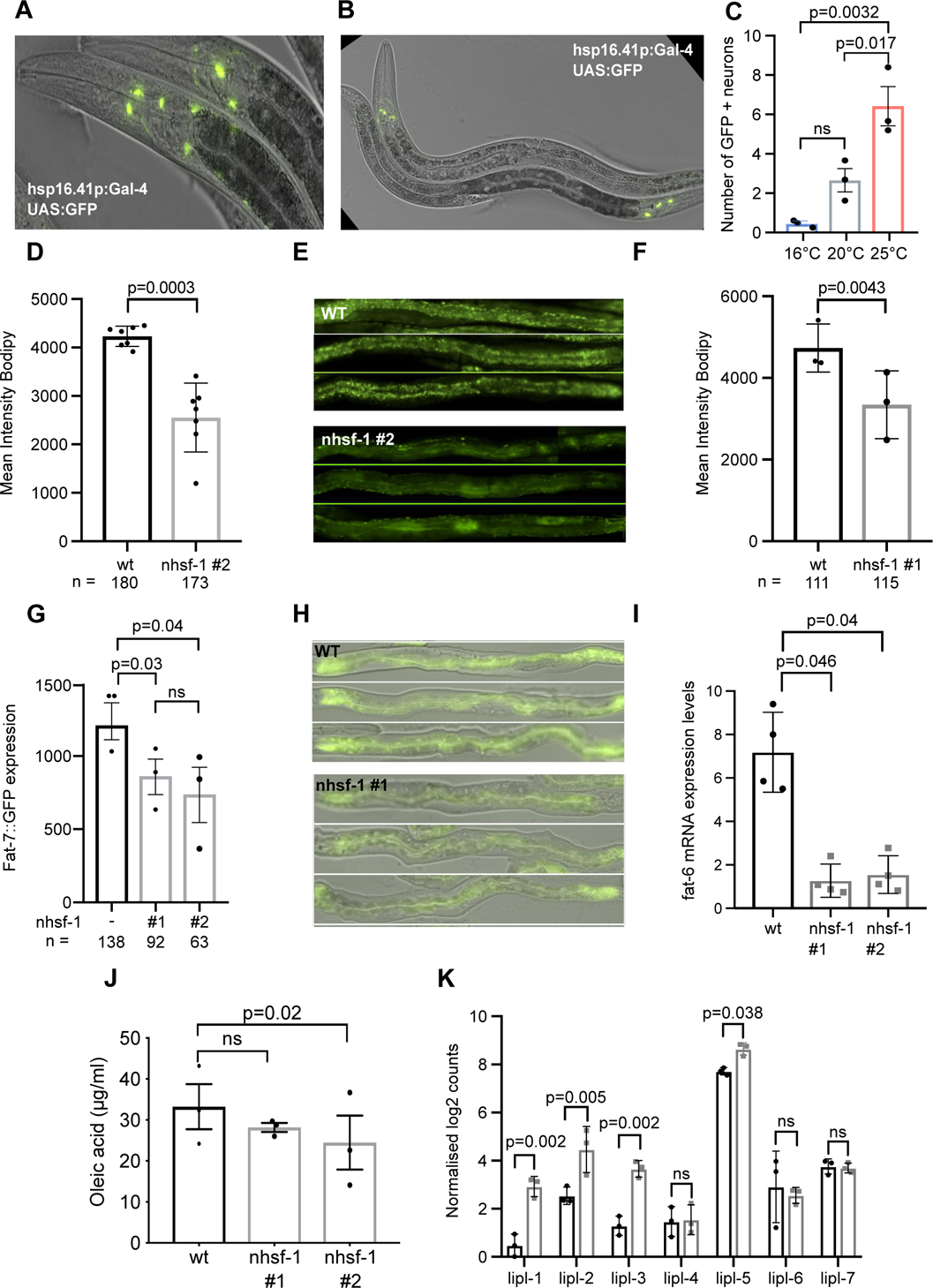
An opposing relationship between neuronal stress and fat metabolism. (**A-B**) An HSF-1-driven promoter is expressed only in anterior neurons. Young adult *hsp16.41p*:Gal4/UAS:GFP animals raised at 20℃. 20X overlay of GFP fluorescence and Nomarski images of GFP expressed exclusively in anterior neuronal cells with axons (in A, arrow head). (**C**) Neuronal expression of an HSF-1 dependent promoter is temperature-sensitive. Quantification of the number of anterior neurons expressing GFP at 15℃, 20℃ and 25℃ in *hsp16.41p*:Gal4/UAS:GFP. The number of GFP + neurons increases by 6.6-fold from 16℃ to 20℃; 16-fold from 16℃ to 25℃; and 2.5-fold from 15℃ to 20℃. Each dot represents a biological replicate, statistics were performed using a one-way ANOVA test. All data are provided in Table S1A. **(D-F)** Wild type and *nhsf-1* over-expression animals from line 1 (**D**) and line 2 (**F**), stained with BODIPY, which intercalates with lipid droplets (LDs). Mean intensity fluorescence levels are reduced by 30% or 40% in *nhsf-1#1) and # 2* respectively relative to wild type animals, indicating that these animals are leaner relative to wild type worms. Statistics was performed using paired t-test on 7 and 3 biological replicates, respectively. (**E**) Images of straightened animals (upper panel, WT; lower panel, *nhsf-1#2* animals) showing GFP fluorescence in the intestine, imaged at 20x magnification. (**G-H**) Ectopic expression of *hsf-1* in neurons results in the decreased activity of a transcriptional *fat-7* reporter, which drives GFP in the intestine of young adult animals, at 20°C. (**G**) GFP fluorescence is decreased by 28% in *nhsf-1# 1* and 38% in *nhsf-1# 2* compared to wild types. One-way ANOVA was applied to paired biological replicates. (n) total numbers of worms analysed are listed below the figure. (**H**) GFP fluorescence and DIC images overlaid, taken at 20X magnification. **(I)** mRNA of *fat-6* quantified by qRTPCR in wild type versus *nhsf-1* animals. Each dot represents an independent biological replicate obtained at 20°C and the bar is the standard error. Statistics was performed using a paired t-test (Sup Table 1E). **(J)** Results from HPLC/MS analysis, showing levels of total fatty acids (FAs) (in μg per ml of fatty acid of free FAs), as summarised in Supplementary Table 5. The FA, Oleic acid, has an 18 acyl-chain with a double bond in position 18 (C18, c9), and is significantly different levels between wild type and *nhsf-1#*2. Each dot represents a paired biological replicate, the error bar corresponds to standard error and the statistics was performed using One-way Anova (see Supplementary Table 1D). **(K)** Normalised log counts obtained by DESEQ2 from an RNA sequencing comparison of wild type and *nhsf-1*#2. The RNA sequencing experiment is described in the methods. All differentially expressed genes are shown in Table 6. Detectable lipase mRNAs levels are present in each of the three replicates. Statistics were performed using one-way ANOVA, which showed that *lipl-1*, *lipl-2* and *lipl-3* are significantly increased in *nhsf-1*#2 (Sup Table 1F). *lipl-1*, *lipl-2, lipl-*3and *lipl-5* were found to be differentially expressed by DEseq2. All DE genes can be found in Sup Table 6.

The striking temperature-dependent increase in *hsp* expression in anterior neurons is opposite to that of *fat-7*. SCD enzymes work by inserting a double bond into the 9^th^ carbon of either palmitic acid (FAT-5) or stearic acid (FAT-6 and FAT-7) (Watts and Ristow, 2017). The FAT-5 desaturase is specific for the synthesis of palmitic acid (16:0), whereas the FAT-6 and FAT-7 desaturases, which share 86% homology at the nucleotide level, mainly desaturate stearic acid (18:0), producing a C18:1 MUFA called oleic acid (C18:1(9z), OA) (Brock et al 2006b; Watts and Browse, 2000) . Consistent with previous reports (Murray et al. 2007; Savory et al, 2011b; Gómez-Orte et al., 2018), we observe that there is a 38-40% decrease in transcript levels of *fat-6* and *fat-7* as the temperature increases from 15℃ to 20℃. The levels of *fat-6* do not decrease further at 25℃ (**Figure S1C**), though *fat-7* declines to 6% of its levels at 15℃ (**Figure S1B, Supp. Table S1B**). Together, these results indicate that the activation of the HSR in anterior neurons has an opposite expression pattern to *fat-6/7* expression in the gut. These observations prompted us to study the potential relationship between neuronal HSR and the expression of enzymes that are known to modulate membrane fluidity.

### Neuronal stress causes reduced oleic acid and fat stores

To explore the role of heat stress responses in neurons, we used transgenic *C. elegans* lines where *hsf-1* expression is driven by the promoter *rab-3*, which is expressed only in neurons and depleted from other tissues (*nhsf-1*) (Douglas et al, 2015; Nonet et al, 1997). We used two integrated *nhsf-1* lines (*nhsf-1*(1) and *nhsf-1*(2) that overexpress *hsf-1* at different levels in head neurons (**Figure S1D, Supp. Table S1C**). Others have previously reported that these animals have increased longevity and heat stress resilience(Douglas et al. 2015) but in addition to these phenotypes, we observed a clear intestine, a phenotype usually associated with decreased fat stores (McKay et al. 2003).This observation indicates that the ectopic expression of *hsf-1*in neurons is potentially linked to fat metabolism in peripheral tissues. *C.elegans* does not have dedicated adipocytes but rather stores fats in organelles called lipid droplets (LDs) in the gut and hypodermis (Watts and Ristow 2017). Electron microscopy and lipidomic analyses have shown that the core of LDs is composed of triglycerides (TAGs), enclosed by a monolayer of phospholipids (PLs) and protein (Zhang et al. 2010; Vrablik and Watts 2013).

Lipid droplets (LDs) can be readily quantified using the fluorescent dye Bodipy (493/503). When Bodipy’s nonpolar structure binds to the neutral lipid components of LDs, it emits a green fluorescence signal with a narrow wavelength (Ashrafi et al, 2005). In young adult nematodes with increasing levels of *hsf-1* in their neurons, we observed a dose-dependent reduction in Bodipy fluorescence of 30% in *nhsf-1(*1) and 40% in *nhsf-1*(2), relative to wild type animals (**Figure 1** **D-F, Supp.Table 1B**). LDs are a vital energy source during periods of low food availability(Watts and Ristow 2017). A potential cause, therefore, for decreased LD levels in *nhsf-1* worms is that the ectopic expression of *nhsf-1* causes feeding to cease, resulting in the depletion of fat stores. However, the expression of *mir-80*, a micro-RNA that is upregulated in starved animals(Vora et al. 2013), is not different in *nhsf-1* young adult animals compared to age-matched, wild type controls (**Figure S1E, Table S1F**). In addition, pharyngeal pumping rates, which when slowed can reduce or preclude a worm’s feeding ability, were also not significantly different in *nhsf-1* nematodes relative to wild type, age-matched controls (**Figure S1F, Supp. Table S1D**). These observations indicate that LD depletion in *nhsf-1* is not due to starvation.

One of the FA components of LDs are C18 MUFAs (Zhang et al. 2010; Vrablik and Watts 2013). Because the *fat-6; fat-7* double mutant show a dramatic decrease in the levels of oleic acid (OA) and LDs (Shi et al, 2013), we assessed the expression levels of Δ9 desaturases in *nhsf-1* worms. We found that an *in vivo* transcriptional *fat-7* reporter (Walker et al. 2011) was reduced in a dose-dependent fashion in the two *nhsf-1* lines by 28 % in *nhsf-1*(1) and by 38% in *nhsf-1*(2) relative to wild type (**Figure 1G, H**). At the transcript level, *fat-6* is also decreased by almost 7-fold in *nhsf-1* worms (**Figure 1I** Supp.Table 1E).

If the observed decrease in the transcriptional output of these enzymes in *nhsf-1* worms was accompanied by reduced enzymatic output, then we would expect to observe changes in vaccenic acid and OA levels. To quantify FA composition in young adult *nhsf-1* worms relative to age-matched, wild type controls, we used chromatography coupled to mass spectrometry (GC-MS) to measure levels of total FAs. This analysis showed that although the composition of most free FAs remained stable in *nhsf-1* lines, the levels of C18:1 OA was significantly reduced by 27% with respect to wild type in the presence of neural *hsf-1* over-expression (**Figure 1J**, **Supp.** Table 1**D and Supp.Table 5J**). These results indicate that ectopic neuronal stress causes the remodeling of LDs, at least in part by compromising the enzymatic output of the Δ9 SCD, FAT-7. It has previously been shown that the loss of a transcriptional activator of *fat-7*, MDT-15 and the loss of *fat-7* itself, can downregulate *hsps* at 15°C (Lee et al. 2019b). To test if reduced *fat-7* levels feedback to further augment stress in neurons at 20°C, we partially knocked down the function of *fat-7*and *fat-6* using RNA interference (RNAi) and tested the effect on the output of UAS/GAL4 driven by the promoter of *hsp16.41.* As shown in **Figure S1G-H, Supp. Table S1E,** reduced Δ9 SCD levels did not change neuronal stress, indicating that above 15°C, Δ9 SCD activity functions only downstream of neuronal stress.

LDs are catabolized by lysosomal lipases, which then recycle the FAs that coat LDs back into the cytosol, where they can be broken down by β-oxidation or recycled to membranes (Watts and Ristow 2017). *C. elegans* contains at least five lysosomal lipases, LIPL-1 to LIPL-5 (O’Rourke and Ruvkun, 2013; Lapierre et al. 2011; Folick, et al. 2015; Ramachandran et al. 2019) which are activated by the acidic environment of the lysosome. Two fasting-induced lipases, LIPL-1 and LIPL-3, belong to the family of Adipocyte-Triglyceride Lipase (ATG)-like Patatin-domain containing lipases. These two lipases localize to the intestine, and their loss is accompanied by an increase in LDs and in overall body fat (O’Rourke and Ruvkun 2013). We observed that the transcripts encoding *lipl-1*, *lipl-2*, *lipl-3* and *lipl-5* are upregulated in *nhsf-1* (Highlighted in **Figure 1K** and **Figure S3E** and **Supp. Table 1F and S3D**), indicating that their upregulation might contribute to the remodeling of LDs in *nhsf-1*.

### 2. Neuronal stress controls fat metabolism by dampening TGF-β/Sma/Mab signaling

Together our data indicate that neuronal stress remotely controls fat remodeling and that that the presence or absence of a molecular signal must allow two distant tissues to communicate. When performing an in-depth phenotypic characterization of *nhsf-1,* we observed two phenotypes that shed light upon the potential nature of such a signal. First, animals with an extrachromosomal array that drives neuronal *hsf-1* (Ex-*nhsf-1*) expression were 20% smaller in size relative to age-matched wild type animals (**Figure S2A, Supp. Table S2A**). Second, *nhsf-1*(2) animals slowed down germline senescence (Luo et al. 2010). When *nhsf-1(*2) animals were cultured in the absence of males, they self-reproduced as expected, but produced significantly fewer progeny than wild types controls in the first three days of reproduction. However, they then continue to produce progeny at a significantly higher rate than wild type animals and for a longer period of time (**Figure S2B-C, Supp. Table S2B**), similarly to mutants that extend germline senescence (Luo et al. 2010)

Based on these phenotypes, we investigated whether *nhsf-1* animals might phenocopy the loss of TGF-β/Sma/Mab signaling pathway for three reasons. First, the loss of DBL-1, the sole Sma/Mab pathway ligand in *C. elegans*, which is related to vertebrate Bone Morphogenetic Protein 10 (BMP) (Morita, Chow, and Ueno 1999),(Suzuki 1998), causes a small (SMA) phenotype. Second, TGF-β/Sma/Mab signalling regulates reproductive span, in parallel to the Insulin/IGF-1 signaling (IIS) and Dietary Restriction pathways (Luo et al. 2009; 2010). And third, mutations in the TGF-β/Sma/Mab pathway cause a lean phenotype, similar to that of the *nhsf-1* phenotype. In addition, mutations in TGF-β/Sma/Mab pathway genes decrease the abundance of LDs (Clark et al. 2018; Yu et al. 2017) as well as the expression of the SCD desaturase genes, *fat-6* and *fat-7* (Yu et al. 2017; Taylor et al., n.d.; Luo et al. 2009)

To test the hypothesis that neuronal stress dampens BMP signaling, we took advantage of the RAD-SMAD reporter, in which multiple, high-affinity SMAD-binding sites are placed upstream of GFP. Others have previously shown that the RAD-SMAD reporter directly and positively responds to BMP signaling (Tian et al. 2010). As shown in **Figure 2A**, wild type worms grown at the standard temperature of 20°C expressed GFP in both hypodermal and intestinal cells during the L4.8 stage. However, in the presence of *nhsf-1*(2), the reporter’s GFP signal is visibly reduced at the same developmental stage (see **Figure 2B**, which shows reduced reporter expression in individual *nhsf-1*(2) worms). Two nuclei types showed a positive RAD-SMAD signal: small (< 8µm) and large (>8µm) nuclei, which most likely correspond to hypodermal and intestinal nuclei, respectively. As shown in **Figure 2C**, the RAD-SMAD signal decreases by 28% in small hypodermal nuclei in the presence of *nhsf-1*(2) compared to wild type levels **(Supp. Table2A)**. A similar pattern was observed in intestinal nuclei, where there was a 53% decrease in the RAD-SMAD signal, relative to wild type levels (**Figure 2D, Supp.Table 2A**). These results indicate that *nhsf-1* partially reduces the activity of the TGF-β/Sma/Mab pathway.

**Figure 2.**
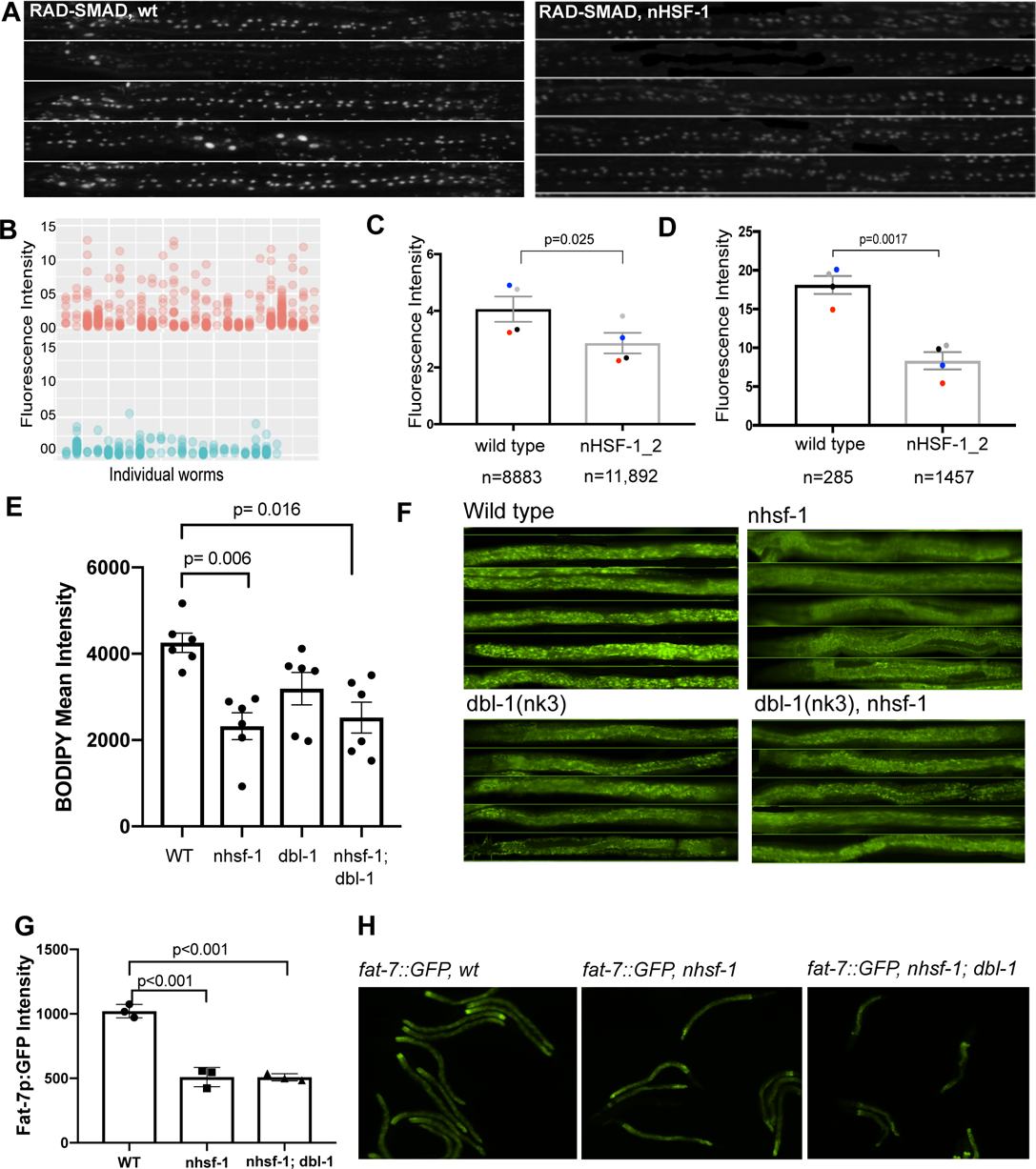
Neuronal stress remodels fat metabolism by modulating BMP signalling. (**A**) Images of worms that expression an in vivo RAD-SMAD reporter that directly and positively responds to SMA signalling (Tian, 2010) (LW2436: jjIs2277 [pCXT51(5*RLR::pes-10p(deleted)::GFP) + LiuFD61(mec-7p::RFP). The reporter can be visualised as a GFP signal in the nuclei of hypodermal (small nuclei, <60um) and intestinal cells (larger nuclei>60um) in wild type L4.8 animals (left panel) or in L4.8 nHSF-1_2 animals (right panel), grown at 20℃. (**B**) Quantification of the RAD-SMAD reporter. Each column shows the intensity of GFP in individual worms from a representative replicate. Red dots represent fluorescent intensity for RAD-SMAD in wild type animals and blue dots represent the intensity of RAD-SMAD in nHSF-1_2 worms. (**C**) Quantification of RAD-SMAD levels in large nuclei, most likely corresponding to intestinal nuclei. There is a significant difference in GFP levels in wild type worms relative to nHSF-1_2 animals (paired t-test) (Sup Table 2A). N values correspond to the number of measured nuclei, across all worms from 4 independent replicates. Paired-replicates have been colour-coded across treatments. (**D**) Quantification of RAD-SMAD levels in small nuclei, most likely corresponding to hypodermal nuclei. There is a significant difference in GFP levels in wild type worms relative to nHSF-1_2 animals (paired t-test) (Sup Table 2A). N values correspond to the number of all measured nuclei, across 4 independent replicates. Paired-replicates have been colour-coded across treatments. (**E**) Genetic epistasis analysis using the following genetic backgrounds: tax-2(p81); dbl-1(nk-3); nHSF-1_2. Each dot corresponds to the average intensity level of Bopipy fluorescence per biological replicate. Statistics, one-way ANOVA. N values (Sup Table 2B). (**F**) Representative images of Bodipy fluorescence for wt, single and double mutants taken at 20x magnification. Scale bar (**G**) Quantification of fat-7p::GFP fluorescence in WT, nhsf-1 and nhsf-1;dbl-1(nk3) genetic backgrounds. Statistics was performed using a one-way ANOVA. (**H**) Representative images of fat-7p::GFP fluorescence in WT, nhsf-1 and nhsf-1;dbl-1(nk3) genetic backgrounds, taken at 20X magnification. All statistics are summarised in Sup Table S2A-B.

To further test for this potential interaction, we performed genetic epistasis experiments. We hypothesized that if *nhsf-1* reduces TGF-β/Sma/Mab signaling, then in the absence of DBL-1, *nhsf-1* should not be able to further reduce LD accumulation. To test for this, we used *dbl-1(nk3),* which carries a large deletion in the coding region of *dbl-1,* and monitored the accumulation of LDs in this mutant via BODIPY staining. As described elsewhere (Clark et al. 2018; Yu et al. 2017) this BMP mutant has 20% lower levels of LD accumulation, relative to wild type animals (**Figure 2E, Supp.Table 2B**). However, in the double *dbl-1(nk3)*/*nhsf-1*(2) mutant, LD accumulation levels are not significantly different relative to those observed in *nhsf-1*. In addition, and as described elsewhere (Luo et al. 2009; Yu et al. 2017) the loss of *dbl-1* function causes a reduction in *fat-7* expression, in line with our observations for *nhsf-1* (**Figure 1**). Our findings reveal that relationship exists between *nhsf-1* and *dbl-1* with regards to *fat-7* expression, because the double mutant is similar to the *nhsf-1* single mutant (**Figure 2** **G-H**). Together, these results suggest that the loss of LD accumulation in *nhsf-1* is caused at least in part, by the loss of TGF-β/Sma/Mab signaling pathway activity. In further support of an epistatic relationship, we observed that the *nhsf-1* transgene when crossed onto a *dbl-1(nk3)* genetic background did not cause the double mutants to be shorter than either single mutant (**Figure S2D**).

### 3. Neuronal TAX-2/TAX-4 expression is essential for the neuronal-stress fat-remodeling phenotype

To identify the neurons that transmit the stress-related signal that results in fat remodelling, we performed a suppressor RNAi screen in *nhsf-1* animals. Because neurons tend to be refractive to RNAi, we combined *nhsf-1* with a loss of function allele in *rrf-3*, which enhances RNAi sensitivity, including in neurons(Simmer et al. 2002) .Animals were fed with bacteria producing RNAi against seven genes that are required for sensory neuron function, including *tax-4*, *ttx-3*, *cat-2*, *tax-6*, *tbh-1*, *tph-1*, *unc-31*. Specifically, we screened for the dampening of *fat-7p:*GFP expression in RNAi-bacteria-fed *nhsf-1* worms. Because *fat-7* expression levels are highly sensitive to dietary variations, the screen was performed using an extra-chromosomal array, which carried the transgene *rab-3*P: *hsf-1* (henceforth called Ex *nhsf-1*). In this experimental setup, due to the incomplete transmission of extrachromosomal arrays, siblings of different genotypes were grown side by side under identical conditions. The effect of RNAi treatment on *fat-7p*:GFP in siblings that inherited the array was compared to those that had not (**Figure S3A-B, Supp. Table S3A**). As expected, in animals fed with an empty vector (EV), demonstrated a significant 37% reduction in the transcriptional output of the *fat-7p: GFP* reporter in *nhsf-1*, compared to wild type siblings. We found that four of the RNAi treatments acts as suppressors of the *fat-7* reduction in *nhsf-1*. However, two of these four were non-specific suppressors because they caused an increase in *fat-7*pGFP in control siblings (Supp. **Table S3A**), the exceptions being *tax-6* and *tax-*4. As the normalized data shows, while *tax-6* RNAi causes a mild suppression, *tax-4* RNAi reverts *fat-7* expression to almost wild type levels (**Figure S3B**). TAX-2 and TAX-4 are respectively, the α and β subunits of a hetero-oligomeric cyclic nucleotide-gated channel that is required for the proper functioning of several sensory neurons and that is also involved in chemosensation, thermotaxis and Dauer formation(Komatsu, Mori, and Rhee 1996; Coburn and Bargmann 1996).

Because RNAi causes only a partial knock-down of gene function and introduces technical variability, we further investigated these results using loss of function mutations. TAX-2/TAX-4 are expressed in 14 sensory neurons in the head, including in the AFD neuron (White, et al 1986; Coburn and Bargmann 1996; Komatsu et al,1996). AFD is the main thermosensory neuron, is required for thermotaxis (Satterlee et al, 2001; Wang et al, 2013; Hobert et al, 1997) and has been shown to key for the *nhsf-1* modulation of the peripheral heat shock responses (Prahlad et al, 2008; Douglas et al. 2015). The DBL-1 ligand is also expressed in several neuronal cell types, including the ventral cord neurons and the AFD neuron (Morita et al, 1999). An obvious potential mechanism would be that neuronal stress directly turns off the expression of DBL-1 ligand in the thermosensory AFD neuron. However, it is unclear if DBL-1 is expressed in AFD as others have argued that it is instead expressed in AVA interneurons (Zhang and Zhang, 2012).

In our RNAi screen, we ruled out the involvement of *ttx-3*; *ttx-3* encodes a LIM-homeodomain protein, which is expressed in the AIY neuron, and is required for thermotaxis (Hobert, 1997). To completely rule out the involvement of the AFD neuron in fat metabolism, we used genetic mutations in *ttx-3* and in *ttx-1*, which encodes a homeodomain-containing protein that is necessary and partially sufficient for AFD development (Pereira et al. 2017). Our results indicate that *nhsf-1* only partially suppresses the activity of a BMP signaling sensor (see **Figure 2**); we therefore reasoned that removing a potentially important source of DBL-1 (the AFD) should further increase the phenotypic penetrance of the *nhsf-1* transgene with regards to *fat-7* expression and body size. However, the tested *ttx-1 (p767) and nhsf-1* (**Figure 3A, C, Supp.Table 3B**) showed a negative interaction in a two-way ANOVA test. These results indicate that the difference between wild type vs *nhsf1* is similar to the difference between *ttx-1* and *ttx-1; nhsf-1*. Likewise, a negative two-way ANOVA interaction was found for *ttx-3 (ks5)* mutants (**Figure S3C, Supp. Table S3B**). It nevertheless remains possible that if *nhsf-1* eliminates *dbl-1* expression only from the AFD. If this was the case, then genetically compromising the AFD function would not enhance the *nhsf-1* phenotype. However, if this were correct, then eliminating the function of the AFD in a wild type background should alter LD accumulation. Our results suggest that this isn’t the case because none of these mutants on their own shows a significant change in *fat-7* promoter-driven GFP expression relative to wild type (**Fig 3** **A and C, Fig S3 C, Supp.Table 3B, Supp. Table S3B**), indicating that the AFD *per se* does not modulate fat remodeling in worms. These results also rule out any involvement of the thermosensory AFD neuron on the modulation of Δ9 SCD expression. They also indicate that the effect of *nhsf-1* on stress resistance, which depends on AFD function (Douglas, et al 2015), is separable from the effect of *nhsf-1* on body fat distribution.

**Figure 3.**
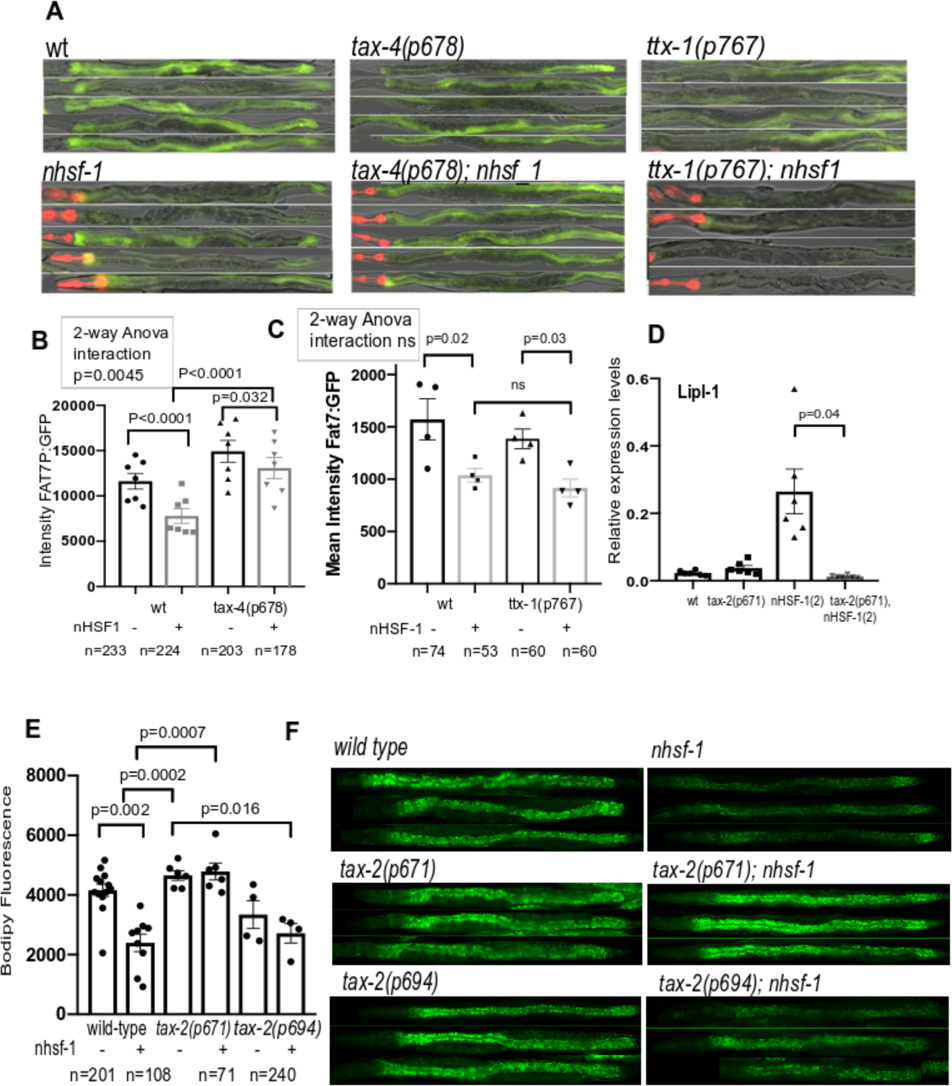
Neuronal cells expressing the cGMP receptors *tax-2/tax-4* are required to modulate the effect of neuronal stress on fat-desaturation in the gut. (**A-C**) Genetic epistasis using *uthEx663[rab-3p::hsf-1; myo-2p::tdtomato]; Is[fat-7p::GFP]* Crossed to *tax-4(p678)* or *ttx-1 (p767).* Siblings were directly compared to monitor the effect of the mutation on the transcriptional output of fat-7p:GFP reporter in a wild type or neuro-Hsf-1 context. GFP is expressed from *fat-7*p:GFP and RFP is expressed from *myo-2p::tdtomato*, a co-injection marker only expressed in NHSF-1 animals and used to distinguish genotypes among sibling. (**A**) 20x fluorescence and brightfield images of representative animals of the genotypes described. (**B-C**) Quantification of the epistasis experiment to determine the interaction between *tax-4* and *nhsf-1*, with respect to the transcriptional output of the *fat-7p*:GFP transgene. Y axis is the mean intensity measured on green channel +/-SEM. Each dot represents a replicate, N values are shown beneath the graph and the gray columns highlight siblings containing the *nhsf-1* array. Two-way ANOVA was performed using GFP intensity values (Sup Table 3A-B). The positive interaction indicates that the difference caused by the presence of nHSF1 is greater in control animals than in *tax-4(p678* (**B**) but not in AFD-deficient animals of the genotype *ttx-1 (p77)* **(C). (D)** Relative expression levels of *lipl-1* mRNA by qRTPCR in wild type, *tax-2(p671),* nHSF-1(2) and *tax-2(p671); nhsf-1*(2) double mutant animals. The levels of *lpl-1* are suppressed in the absence of *tax-2* function. Each dot represents a biological replicate and the bars correspond to the standard error. Statistics was performed using One-way ANOVA (Sup Table 3C). (**E-F**) Epistasis analysis to determine the effect of *tax-2* on NHSF-1_2 with respect to lipid droplet accumulation. The genotype of the strains compared are: *nhsf-1*(2)*; tax-2(p671),* which was compared to *nhsf-1*(2); *tax-2(p694)* and *nhsf-1*(2); *tax-2(p694),* Sup Table 3D **(E)** Y axis shows the average across replicates of mean BODIPY expression, where each dot is one biological replicate, the bar is the standard error, and statistics were performed using one-way ANOVA (**E**) 20x fluorescence images showing fat deposits in representative animals of the genotypes highlighted. The green channel shows the levels of Bodipy intercalated in lipid droplets present in the gut and the hypodermis. All statistics have been summarised in Sup Table 3.

To corroborate the suppressive effect of *tax-4* RNAi, we used a loss of function allele, *tax-4(p674)* (Satterlee, et al 2014). As shown (**Figure 3** **A, B)**, the presence of *nhsf-1* significantly decreases the expression of *fat-7p: gfp* by 32% compared to wild types. In the absence of *tax-4,* GFP expression was not significantly altered (**Supp**. **Table 3A)**. However, the *tax-4(p68)* mutation partially suppressed the effect of *nhsf-1* on *fat-7* expression, increasing *fat-7p: gfp* expression to almost wild type levels (**Figure 3A-B, Supp.Table 3A**). A two-way ANOVA test shows a positive interaction, indicating that the difference between sibling pairs is higher for the control pair than for animals bearing the *tax-4(p678)* mutation. Together, these results indicate that eliminating *tax-4* function partially suppresses the inter-tissue effect of *nhsf-1* with respect to *fat-7* expression in the gut. A further analysis indicated that *tax-4* does not suppress the small phenotype of *nhsf-*1 (**Figure S3D, Supp. Table S3CD)**, separating the regulation of body size by *nhsf-1* from its regulation of fat metabolism.

In addition to effecting Δ9 SCDs, *nhsf-1* also transcriptionally upregulates lipases, including *lipl-1* by 11-fold (**Figure 3D** **and Supp.Table 3C**) and *lipl-3* by 6-fold (**Figure S3E and Supp. Table S3B**). We therefore used *tax-2(p671)*, a missense mutation in a conserved region of the predicted pore region of the α subunit of the cGMP channel (Coburn and Bargmann, 1996), to test for a potential interaction with lipases. Our findings show that the *tax-2(p671)* mutation suppressed LIPL levels; in the compound *tax-2(p671); nhsf1* mutant, *lipl-1* and *lipl-3* expression levels returned to near wild-type levels. If transcriptional suppression of fat metabolizing enzymes is also accompanied by a change in FAs, then we would expect the elimination of cGMP function to suppress the *nhsf-1* phenotype of depleted LDs. As observed before (**Figure 3E-F, Supp.Table 3D**), *nhsf-1* caused a significant 63% reduction in LDs relative to wild type. *tax-2(p671*) did not significantly alter LD levels in wild type animals, but rescued LD levels in *nhsf-1* (2). Together, our data shows that eliminating the function of TAX2/TAX4 can suppress the expression of enzymes that are required for LD homeostasis with downstream consequences for fat stores.

What might the underlying mechanism for this suppression be? It is possible that cGMP-gated channels help to detect or to transduce the HSF-1 dependent transcriptional response in neuronal cells. To test this hypothesis, we examined the ability of neurons to activate an HSF-1 dependent target, *hsp16.41*, in the absence of *tax-2* function. As shown in **Figure S3F-G**, in *tax-2(p671*) animals that lack a functional cGMP channel, some unidentified neurons in the head region of GAL4/UAS animals can still activate *hsp16.41*. From this observation, we conclude that the cGMP receptor does not participate in detecting or transducing stress responses *per se* in neuronal cells. We support the view that HSF-1 activation in some of TAX-2/4-expressing neurons might directly or indirectly repress the TGF-β/Sma/Mab pathway. Loss of TAX-2/4 might disrupt the functioning of key neurons, disabling the ability of the HSR to influence TGF-β/Sma/Mab signaling.

TAX-2 and TAX-4 are expressed by a subset of 14 overlapping sensory neurons (White et al, 1986; Coburn et al, 1996; Komatsu et al, 1996). *Tax-2(p694)* is an allele with a deletion in the promoter region of *tax-2* that disrupts its expression only in AQR, ASE, AFD, BAG PQR and URX neurons (Coburn and Bargmann 1996; Bretscher et al. 2011). We find that that this allele does not suppress the fat accumulation phenotype of *nhsf-1* (**Figure 3E-F, Supp.Table 3D**). This indicates that none of these neurons is involved in modulating fat remodeling, and that all or possibly some of the remaining neurons that express the cGMP channel, including ASG, ASJ, ASK, AWB, ASI, AWC must be responsible for control of body fat in response to neuronal stress. **Supp.** **Figure 4** summarizes a model that explains how the activation of HSF-1 in cGMP-expressing sensory neurons either directly or indirectly dampens the TGF-β/Sma/Mab signaling pathway, causing a decrease in fat desaturation and LD stores in peripheral tissues.

**Figure 4.**
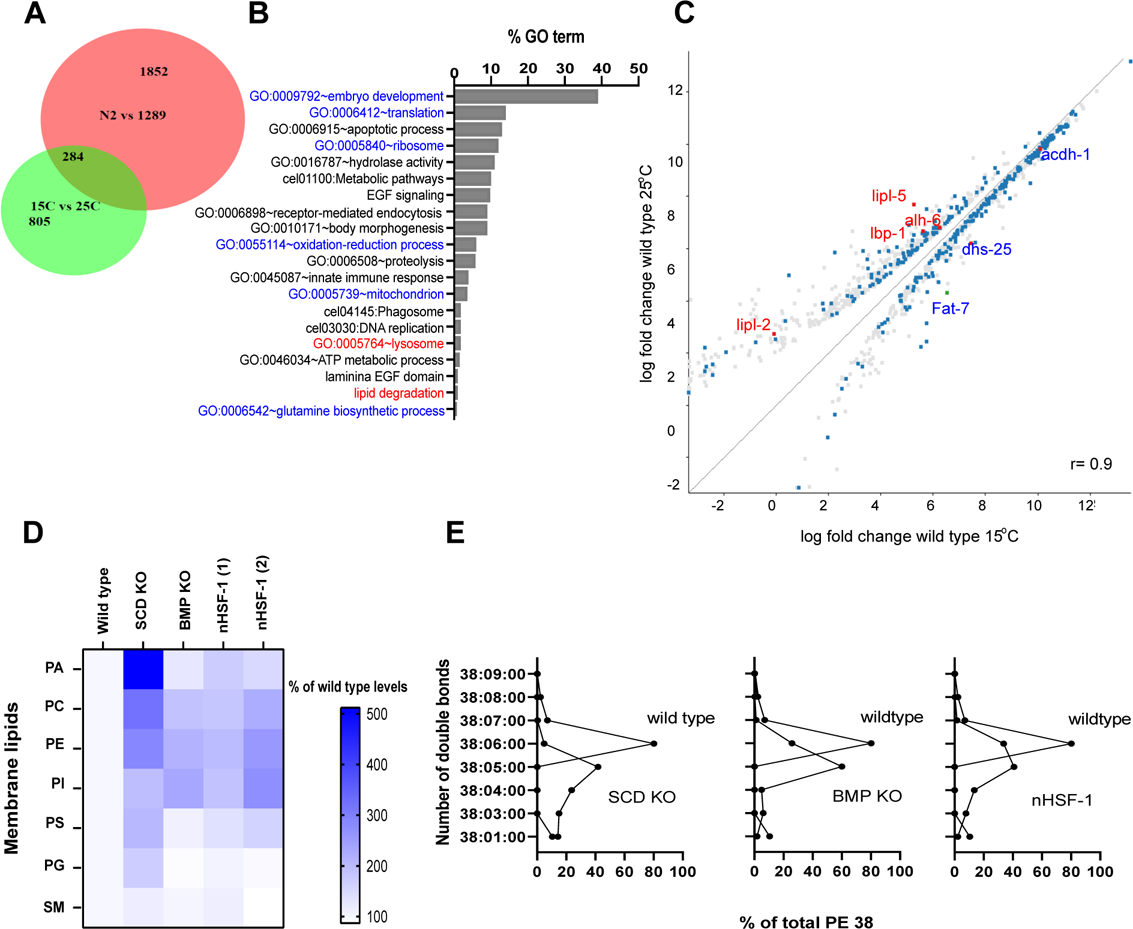
Ectopic expression of *hsf-1* in neurons is sufficient to decrease the fluidity of membranes. (**A**) Overlap between genes differentially expressed between 25°C and 15°C (in the green circle) (raw data taken Lopez Ortis, 2018 and re analysed using Desq-2 using a p<0.05 cut-off, which gave a total of 1,089 genes) and genes differentially expressed between wild type and nHSF-1(2) (in the red circle) (Desq-2 using a cut-off p<0.005, which gave 2,136 genes). A hypergeometric distribution was employed to determine the probability of overlap when using different filtering criteria (0, 0.05, 1, 1.5 or 2 log2-fold change. The overlap was found to be significantly different from chance using any of the mentioned filtering criteria (Sup Table 4B). (**B**) % of enrichment in gene ontologies (GO) found using DAVID of the 284 genes that overlap between both experiments. GOs are highlighted if most genes within that class were downregulated (blue) or upregulated (red) in both nHSF-1(2) and in worms grown at 25°C. (**C**) Scatter plot highlights the correlation (where x and y are shown in the same scale) of the quantified values (in log scale) for wild type animals grown at 25°C versus wild type animals grown at 15°C. The probes highlighted in blue correspond to the overlap between the two experiments (15°C vs 25°C from Lopez Ortiz, 2018 and the comparison between wild types and nHSF-1s). Overlapping probes that are related to lipid metabolism are highlighted in red. (**D**) Heat map showing the abundance of fatty acids found in glycerophospholipids (in ng per ng of DNA), normalised to the abundance of each specific FA in wild type worms. The scale in shown on the right. Although there is a trend for most FAs to increase in the mutants, only PC, PI and PE were statistically significant (Sup Table 4A); PA: phosphatidic acid; PC: phosphatidylcholine; PE: phosphatydilethanolamine; PI: phosphatidyl inositol; PS: phosphatidyl serine; PG: phosphatidyl glycerol; SM: sphingomyelin. (**E**) Relationship between the quantity (in ng/ng DNA) of acyl chain (carbon number related to the fatty acid composition in position *sn-1* and *sn-2* in glycerophosphoplids) for the specified number of double bonds (from 0, saturated to 9, poly-unsaturated) in PE 38 (phosphatidylethanolamine with an acyl chain of 38 carbons). The X axis represents the % of each particular species with regards to the total amount of FA in each mutant. Other examples are shown in Figure S6 and S7. (**D-E**) BMP KO: *dbl-1(nk3)* and SCD KO: steroyl-CoA desaturase: *fat-6 (tm33)1; fat-7 (wa36)*.

### Overexpression of *hsf-1* expression in neurons is sufficient to fine-tune membrane fluidity

At 25℃, *fat-7* expression in the gut is very low, while the number of neurons mounting a stress response is upregulated (**Figure 1** **and S1**). Because the ectopic expression of *hsf-1* in neurons is sufficient to reduce *fat-7* expression in the gut, we propose a model in which sensory neurons in the head, work as a thermostat that detects small increases in temperature to turn on a program that adjusts the fluidity of membranes to better adapt to warmer temperatures (illustrated in **Figure S5)**. Others have already shown that *nhsf-1* causes animals to be more resilient to heat (Douglas et al. 2015). One of the predictions that can be derived from this model is that *nhsf-1* animals have a constitutively active thermostat. If this is correct, then at least some of the transcriptional responses that animals mount when raised at warmer temperatures, should be shared by *nhsf-1* worms.

We identified differentially expressed (DE) genes between *nhsf-1* (2) and wild types by DEseq2 (see materials and methods) and found 2,136 DE genes. Among the gene ontology (GO) groups that characterize the genes upregulated by more than one log2-fold change, include defense response and innate immunity genes, genes required for the formation of ribonuclear proteins, genes involved in reproduction, and genes involved in lysosome function. These data were compared to DE genes from published sources that compared the transcriptomes of animals grown at 15℃ with animals grown at 25℃, under a standard OP50 *E coli* diet (Gómez-Orte et al. 2018). To determine the probability of overlap between the two datasets, we employed a hypergeometric distribution using different filtering criteria (0, 0.05, 1, 1.5 or 2 log2-fold change, (**Table 7**). Using any of these, the overlap was found to be significantly different from an overlap expected by chance. The Venn diagram in **Figure 4A**, shows that the two sets of DE genes overlap by about 10%. **Figure 4B** shows the gene ontology (GO) categories of the overlapping genes. Among the GOs that are common to animals grown at 15℃ and that are down-regulated in *nhsf-1* animals, are: translation control, ribosome formation, amino acid catabolism, and mitochondria. Interestingly, a common GO category that is upregulated both at 25℃ and in *nhsf-1* animals includes genes required for lipid degradation and that are present in the lysosome (see **Figure 4C**), such as *lipl-2* and *lipl-5*. In addition, the fat desaturase, *Fat-7*, is highly downregulated in animals raised at 25℃ (**Figure 4C**, blue) and in animals overexpressing *nhsf-1* (**Figure 1**). Together, these data indicate that *nhsf-1* might regulate a subset of genes that are required to regulate the synthesis and mobilization of MUFAs.

Lipid metabolism lies at the heart of HVA. To ensure membrane fluidity is maintained at warmer temperatures, PLs in the membranes have lower unsaturation levels in FA chains (Sinensky, et al 1974, Cossins and Prosser, 1978). If *nhsf-1* animals were better suited to warmer temperatures, then we would expect the FA composition of glycerophospholipids to follow a similar rule. To test this idea, we performed liquid chromatography-mass spectrometry to quantify the levels of PLs that make up the bulk of the plasma membrane in animals raised at the standard 20℃ and their FA composition. **Figure 4D** shows a heat map of normalized values of different lipids, quantified in four different genotypes, including in the SCD double mutant *fat-6 (tm33)1; fat-7 (wa36);* in the BMP mutant, *dbl-1(nk3)*; and in the two *nhsf-1*lines. We found that the quantity of phosphatidylcholine (PC), phosphatidylinositol (PI), and phosphatidylethanolamine (PE) were significantly changed in all mutants relative to wild type (**Supp. Table 8**), suggesting that the decrease in LDs is accompanied by an increase in specific membrane PLs.

To examine the acyl chain composition of the glycerophosphoplids by determining the number of double bonds for FAs in position *sn-1* and *sn-2*. **Figure 4E** shows the specified number of double bonds (from 0 = saturated, to 9 = poly-unsaturated) for one example, specifically: PE 38 (phosphatidylethanolamine with an acyl chain of 38 carbons). All genotypes phenocopy each other in that the number of double bonds decreases with respect to wild type animals. **Table 8** show other examples of FAs with a reduced number of double bonds. These results suggest that the membranes of *nhsf-1* worms, and those of the other two mutants that have reduced levels of OA production *(fat-6; fat7* and *dbl-1*), are consequently less fluid and can better adjust to warmer temperatures, even though these animals were grown at the permissive temperature of 20℃. Together, our results indicate that the over expression of *hsf-1* in neurons coordinates an organismal response to decrease the production and storage of MUFAs. They show that *nhsf-1* mutants phenocopy SCD and BMP mutants by decreasing fat stores and by remodeling the composition and fluidity of the plasma membrane.

## Discussion

Our results show that the ectopic activation of the heat shock responses in neurons causes LDs to be depleted from fat store cells, and the widespread remodeling of the abundance and composition of membrane PLs, in a manner that would support HVA to high temperatures. We identify TAX2/4-dependent sensory neurons as being important for this response, and a ligand of the BMP family as being the key signal that must be decreased to communicate this response from the brain to the gut. We propose that neuronal stress in key TAX2/4 neurons acts as a thermosensor that enables heat adaptation at the organismal level by regulating peripheral membrane and protein homeostasis.

The biological thermostat is a basic model of thermoregulation that is used by many species Thermostats generally have three components: (1) afferent thermosensation; (2) central integration in the brain; and (3) an effector output that causes negative feedback. Mammals use thermosensing-TRP channels to sense heat (Madden and Morrison, 2019) however, but this TRP subtype is not present in the *C. elegans* genome (Xiao and Xu, 2009). Nevertheless, *C.elegans* does have thermo-sensing neurons, such as the AFD and the ASJ neurons, in which heat activates a cGMP pathway that opens the TAX-2/TAX-4 ion channel to increase calcium currents. However, the primary mechanism by which temperature is sensed by TAX-2/TAX-4 expressing neurons remains elusive. Here, we report an alternative potential thermostat that involves the activation of the heat shock response in TAX-2/TAX-4 expressing neurons. The TAX-2/TAX-4 cGMP receptor is expressed in 14 sensory neurons. Our analysis of the allele of *tax-2(p694)* indicates that the neurons that are potentially important in mediating the effects of neuronal stress on fat metabolism include: ASG, ASJ, ASK, AWB, AWC, and ASI. Of these neurons, the light and pheromone-sensing neuron ASJ is a candidate of particular interest as it is heat-activated (Ohta et al. 2014) (Zhang et al. 2018) and suppresses the protective effects of cold acclimation to cold resistance(Ohta et al. 2014). Indeed, ASJ shortens lifespan at 15°C by inhibiting the metabolic transcription factor DAF-16/FOXO in the intestine, which is also known to regulate *fat-7* activity (Zhang et al. 2018). It would be interesting to determine whether suppressing TAX-2/TAX-4 exclusively in the ASJ could rescue *fat-7* expression levels.

In addition to optimizing the fluidity of membranes in warmer temperatures, previous work has shown that *nhsf-1* animals are better prepared for surviving heat stress as they can potentiate the expression of HSPs across tissues (Douglas et al. 2015). Our findings show that both responses are separable because while loss of the thermosensory circuit disrupts *nhsf-1*-dependent thermotolerance, but it does not alter the *nhsf-1* lean phenotype. Together, these experiments suggest that higher temperatures might turn-on a thermosensor in wild type animals, to coordinate multiple adaptive responses that help animals thrive in a warmer environment.

A cell autonomous sensor, the Acyl-coA dehydrogenase *acdh-11*, has previously been reported to be upregulated on heat acclimation (at 25°C) and to downregulate cold-responsive *fat-7* by sequestering C11/C12 fatty acids that would otherwise bind to and activate NHR-49, a transcriptional regulator of *fat-7* (Ma et al. 2015). Our RNAseq analysis does not reveal a change in the expression of *acdh-11* in *nhsf-1* animals, so it is possible that both systems act independently. Future work should aim to untangle how these two systems interact to ensure HVA.

These findings can also inform our understanding of heat adaptive mechanisms in vertebrates. Recent mammalian studies suggest that FA desaturation patterns correlate with latitude in mammals that inhabit different environments, a trend that is particularly clear for aquatic and semi-aquatic mammals (Guerrero and Rogers, 2019). Therefore, understanding the regulation of membrane fluidity is relevant to understanding how thermoregulation occurs in some endotherms. The control of SCDs is also increasingly recognized as relevant to human pandemics, such as obesity and metabolic syndrome (Sampath and Ntambi, 2011). It is possible that some of the neuronal circuits and molecular players we describe here have been co-opted to modulate fat metabolism in mammals with BMP signaling being a good example of this. And finally, the ER stress response is highly relevant in mammals. Specifically, ER stress in Pomc hypothalamic neurons improves glucose utilization and insulin sensitivity protecting mammals from diet induced obesity (Williams et al. 2014). It is not known if the activation of HSR in neurons can also have systemic consequences for health in mammals, but this would be a worthwhile avenue to explore.

## Acknowledgements

We would like to acknowledge Boo Virk, for helping with experiments and editing figures; Cindy Voisine, Michael Witting, Jon Houseley, Len Stephens, Rebeca Aldunate and Jane Alfred for feedback on the manuscript. We are also thankful to Andy Dillin and Amy Walker for reagents; Mario de Bono and Julie Ahringer for helpful discussions. We also acknowledge the help of Babraham Institute Facilities and CGC for providing strains.

## 1-General methods

### C. elegans maintenance

Nematodes were grown on NGM plates seeded with *Escherichia coli* OP50 strain at 20**°**C unless otherwise stated, according to standard methods (Brenner 1974).

### C. elegans strains

The full list of *C. elegans* strains used and generated for the purpose of this study can be found in supplemental table A. We noticed that the strain AGD1289 would sometimes lose its clear intestine phenotype when grown for more than a month, so we used freshly thawed AGD1289 worms for less than a month.

### Worm synchronization

Worms were age-synchronized either by egg-lay in a 2h period or by treatment with alkaline hypochlorite solution, according to standard procedures (Stiernagle 2005) for experiments that required large amounts of synchronized animals (such as the Bodipy staining, RNA-seq and lipidomics experiments). For bulk qRT-PCR experiments and for microscopy-based quantification of fluorescent reporters at L4.8 or L4.9 stages, we used worms synchronized by egg-laying grown in parallel at different temperatures. Worms were harvested 97h after egg-laying at 16**°**C, 42h after egg-laying at 25**°**C, and L4.8 or L4.9 worms were picked from a mixed population grown at 20**°**C. The L4 sub-stages were assessed according to the morphology of the vulva, as described in Mok et al, 2015.

### Experimental design

Each experiment was performed in at least three biological triplicates. In each biological replicate, each condition comprised at least 30 individual animals in microscopy experiments (bodipy quantification, fluorescent reporter quantification), 15-20 animals in confocal experiments (RAD-SMAD reporter quantification), 15-30 animals in qRT-PCR experiments, 10 animals for pharyngeal assay, and 20 animals for fertility assay.

### Pharyngeal pumping assay

Synchronized young adult animals from each genotype were singled out and assayed for pharyngeal pumping at the young adult stage. Experiments were done in triplicate with at least 10 worms per condition. Pharyngeal pumping movements were followed for 30s, under a dissection microscope, and each animal was scored at least twice.

### Fertility assay

About 20 worms per condition were singled out in 12 well plates seeded with 50µL OP50 at the L4 stage. Each day, all animals were passed onto new 12 well plates. The F1 progeny laid by each individual worm was scored 2-3 days after the P0 parent had been transferred to the well, when the F1s were either L4 or adults. For ease of scoring, the 12 well plate was left on ice for a few minutes, until the animals were immobilized.

### DNA lysate preparation and PCR genotyping

Between 10 to100 worms were picked into 10 µL of Worm Lysis Buffer (50mM KCl, 10mM Tris (pH 8.3), 2.5mM MgCl2, 0.45% NP40, 0.45% Tween-20, 0.01% Gelatin). Tubes were freeze-thawed once before 1 µL 01mg/mL of proteinase K was added to each tube. Worms were lysed and their genomic DNA was released by heating tubes to 65**°**C for 60-90 minutes. Proteinase K was inactivated by heating to 95**°**C for 15 min. Commonly, 1 µL DNA lysate was added to each PCR reaction. All PCR genotyping reactions were performed with Taq DNA polymerase (New England Biolabs #M0267L) with Thermopol buffer according to manufacturer instructions. The list of PCR primers used for genotyping can be found in supplemental table B.

### RNAi

The RNAi suppressor screen was designed to look for increase of *fat-7*p::GFP levels in animals carrying an overexpression of *hsf-1* in neurons. As neurons tend to be refractive to RNAi, the screen was performed in the MOC201 strain carrying a loss of function allele in *rrf-3(pk1426)*, which enhances RNAi sensitivity, including in neurons (Simmer et al. 2002). Because *fat-7* expression levels are highly sensitive to dietary variations, the screen was performed using an extra-chromosomal array, which carried the transgene *rab-3*P: *hsf-1* (henceforth called Ex *nhsf-1*). In this experimental setup, due to the incomplete transmission of extrachromosomal arrays, siblings of different genotypes were grown side by side under identical conditions. The effect of RNAi treatment on *fat-7p*:GFP in siblings that inherited the array was compared to those that had not. RNAi-mediated knock down of candidate genes involved in neuronal functioning was performed using clones from Dr Julie Ahringer’s RNAi library, including *tax-4*, *ttx-3*, *cat-2*, *tax-6*, *tbh-1*, *tph-1*, *unc-31*. Bacterial cultures were grown overnight in LB with 100 µg/mL Carbenicillin and induced with 1 mM IPTG for 2 h. RNAi seeded plates were left for 48h to dry at room temperature. Animals were initially grown on OP50 plates and were individually transferred at the L4.8 stage (Mok et al, 2015) onto RNAi plates. About 50 extrachromosomal overexpressing neuronal *hsf-1* worms and non extra-chromosomal worms from the same background were transferred onto the same RNAi plate. Animals were transferred onto fresh RNAi plates at day 2 of adulthood. At day 3 of adulthood, the animals were mounted, and *fat-7*p::GFP fluorescence was monitored. We tried to image about 40 extrachromosomal carrying neuronal *hsf-1* overexpression array, and 40 non extra-chromosomal siblings. At least 3 biological replicates of each experiment were performed.

### qRT-PCR on bulk worm samples

To monitor steady-state mRNA levels on bulk samples of worms, we used the Power SYBR® Green Cells-to-C_T_™ kit and hand-picked a pool of about 15-25 animals in 10 µL Lysis buffer. Reverse transcription was performed using the 2X RT buffer from the Power SYBR® Green Cells-to-C_T_™ kit according to manufacturer instructions and cDNA was diluted either 1:4 or 1:5. Each qRT-PCR reaction contained 1.5 µL of primer mix forward and reverse at 1.6 µM each, 3.5 µL of nuclease free water, 6 µL of 2X Platinum^®^ SYBR^®^ Green qPCR Supermix-UDG with ROX (ref 11744-500) and 1 µL of diluted cDNA. The list of PCR primers used for qRT-PCR can be found in supplemental table B. The PCR efficiency was calculated for each couple of primer by running a standard curve on a dilution series. Validated couples of primers had a PCR efficiency between 90 and 113% with R^2^>0.97 (supplemental table B).

### 2-Lipidomics

#### Worm harvesting for lipidomics

Worms of every genotype were synchronized using hypochlorite treatment according to standard procedures (Stiegerale 2005). Young adult animals were grown at 20**°**C and harvested at 50-52h post L1 plating for N2, MOC141 and NU3, at 54h for AGD1289, and at 70h for BX156, as they were developmentally delayed. One 9 cm NGM plate containing either 1000 young adults for total fatty acids lipidomics, or 500 worms for phospholipids analysis was harvested for each genotype. Worms were washed with 15 mL of M9 buffer at least 3 times. Most of the supernatant was removed and the pellets with collected worms were frozen at −80**°**C in low-protein binding Eppendorf tubes, before being processed for lipidomics analysis.

#### GC-MS Analysis of Fatty acids

16-35 mg of *C. elegans* were extracted by adding 1 mL chloroform/methanol (2:1, v:v, containing 0.01 % BHT and 10 µg of C9:0 and C13:0, respectively as internal standard) and processed using an ultrasound sonotrode for 30 sec at 40 Hz (type UW 2070, Bandelin). Afterwards, 0.5 mL water was added to each tube and each sample was shaken vigorously for 1 min. Next, the extract was centrifuged for 5 min at 3000 rpm. The chloroform layer was transferred into a new vial and the solvent was removed with a gentle stream of nitrogen. The residue was resolved in 200 µl tetrahydrofuran. 400 µl methanolic base 0.5 M (Acros Organics) was added. After 1 min of rigorous shaking, the sample was heated for 15 min at 80°C. Afterward, 200 µL water and 200 µl hexane was added. After another minute of vigorous shaking, the sample was centrifuged for 1 min at 1500 rpm. The hexane layer was transferred into a new vial. 1 µL sample was injected into the gas chromatograph coupled to the mass spectrometer (GC-MS 40, Shimadzu) with a split of 5 and injection temperature of 260°C. The separation was performed with a Zebron ZB-5MSplus column (30 m x 0.25 mm x 0.25 µm) (Phenomenex) with helium as carrier gas: 35°C was held for 2 min. Then the temperature was raised by 10°C per minute to 140°C, which was held for 10 min. Afterwards, the temperature was raised to 240°C at a rate of 2°C per minute. 240°C was held for 10.5 min. Quantification was performed using the GCMSSolution Version 4.30 (Shimadzu).

#### Phospholipids analysis

The worms were homogenized using a Precellys evolution with a cryolys unit to keep the sample frozen during homogenization (Bertin Technologies, France). Precellys bead-beating tubes with reinforced walls for hard tissue (CRK28-R) were used with 3 cycles of 7200 rpm at 45 seconds. The homogenized sample was then transferred to a glass tube containing Chloroform/Methanol for lipid extraction using Folch method (Folch, J. *et. al*. J. Biol Chem 1957). Phospholipids were separated using a Cogent HPLC column (150 × 2.1 mm, 4 µm particle size) placed on a Shimadzu XR (Shimadzu, Kyoto, Japan) using the conditions described in Zhuang, X. *et. al*. The phospholipids were then detected using an Orbitrap Elite mass spectrometer in full scan mode with a mass range of 200-1000 *m/z* at a target resolution of 240,000 (FWHM at *m/z* 400). Data were analysed using Lipid Data Analyzer (2.6.0–2) software (Hartler, J. et.at Nat. Methods 2017).

#### 3-Fat content analysis using bodipy

##### Bodipy protocol

We used bodipy 493/503 (Thermo Fisher Scientific D3922) to stain neutral lipids in fixed animals. We adapted the protocol developed by Klapper and colleagues (Klaper et al. J Lipid Res 2011) to fix worms using 60% isopropanol. About 1000 worms per genotype were synchronized by hypochlorite treatment (Stiernagle, 2005). Animals were collected and washed at least 3 times in M9 buffer. Worms were then fixed for 5 minutes in 60% isopropanol in 1.5 mL protein-low bind Eppendorf tubes, with occasional inversion of the tubes. As fixation with isopropanol was sometimes variable, we also tried fixation with cold methanol for 10 minutes, the rest of the protocol remaining identical After fixation, we let the worms settle by gravity and washed them once more with M9. The M9 supernatant was removed, leaving approximately 50 µL. The tubes were frozen and thawed twice and then incubated with 500 µL of bodipy 493/503 (diluted in M9 at 1µg/mL) at room temperature for 1h on a rotator. After 1h, the worms were washed twice with 1 mL M9 solution containing 0.01% triton. We kept the samples at 4°C and imaged them either the same day or the next day but not more than 2-3 days after collection.

##### Mounting of fixed animals

For mounting, we used a mouth pipette with a glass capillary to remove all liquid in the Eppendorf tube. About 8-10 µL of Vectashield^®^ antifade mounting media without DAPI (Vector Laboratory 94010) was added and worms were mounted for imaging on a 2% agarose pad using a 18×18 mm glass coverslip.

##### 4-Microscopy

All worms imaged were mounted on a 2% agarose pad. For imaging of live worms, animals were paralyzed in 3mM Levamisole diluted in M9. Fluorescence exposure was identical across all conditions of the same experiment. To image intestinal levels of *fat-7p::GFP* fluorescence in live worms and BODIPY 493/503 fluorescence in fixed worms, we used a Nikon Ti Eclipse fluorescent microscope at objective 20X. For imaging GFP-positive neurons in PS7171 and PS7167, we used a Nikon fluorescent stereomicroscope SMZ18, as it was easier to capture neurons in 2D from live animals. PS7171 and PS7167 were synchronized at L4.8 stage (Mok at al., 2015) in this experiment. LW2436 and MOC229 worms were imaged on a Nikon A1R confocal microscope at 20X objective to determine the fluorescence levels of the RAD-SMAD reporter. The Z-stacks taken were then processed by deconvolution, and stitched together.

##### Image Analysis

In order to straighten and quantify fluorescence from acquired *C. elegans* images, we used the FIJI/ImageJ worklow called “Worm-align” that allows to generate single-or multi-channel montage images of aligned worms from selected animals in the raw image. The output of “Worm-align” was then imported and run through a CellProfiler pipeline called “Worm_CP” for fluorescence quantification of the animals of interest selected with “Worm-align”. Both “Worm-align” and “Worm_CP” pipelines are available and described in Okkenhaug et al. JoVE, 2019, in revision.

##### 5-RNA sequencing

###### RNA-sequencing library preparation

RNA was extracted by standard Trizol extraction techniques. Libraries were made using either NEBNext mRNA Second Strand Synthesis Module (E6111) followed by NEBNext Ultra II DNA Library Prep Kit for Illumina (NEB-E7645) or NEBNext Ultra II Directional RNA Library Prep Kit for Illumina (E7760) with the NEBNext Poly(A) mRNA Magnetic Isolation Module (NE7490) as per manufacturers protocols using half reactions. 13 Cycles of amplification was used for library enrichment; quality and size distribution of the the libraries was ascertained by running on a Bioanalyzer High Sensitivity DNA Chip (Agilent 5067-4626) and concentration was determined using KAPA Library Quantification Kit (KK4824). Libraries were sequenced on an Illumina HiSeq 2500 system by the Babraham Sequencing Facility.

###### RNA-sequencing analysis

The FASTQ files were quality trimmed with Trim Galore v0.4.4 (https://github.com/FelixKrueger/TrimGalore), used in conjunction with Cutadapt v1.15 (<u>DOI:10.14806/ej.17.1.200</u>), and then mapped to the *C. elegans* reference genome WBcel235 using HISAT2 v2.1.0 (Kim et al), in either single- or paired-end mode depending on the sequencing protocol followed. To improve the mapping efficiency across splice junctions, HISAT2 took as additional input the list of WBcel235.75 known splice sites. RNA-seq QC and analysis was performed on the mapped reads using the genome browser SeqMonk v1.45.4 (https://www.bioinformatics.babraham.ac.uk/projects/seqmonk/). To identify differentially expressed genes, the number of reads positioned over exons were first calculated for every gene using the program’s “RNA-seq Quantitation Pipeline”. Differentially expressed genes were subsequently called by DESeq2 v1.22.2 (Love et al.,), launched from SeqMonk with default settings. Differentially expressed genes were defined by having an adjusted P value cutoff <0.05 after multiple-testing correction. The gene expression Principal component analysis when comparing *nhsf-1* and wild type showed that the transcriptomes formed clusters according to their molecular subtypes, indicating high quality and consistent homogeneity of transcriptomes.

---

**Supplementary Figure 1.**
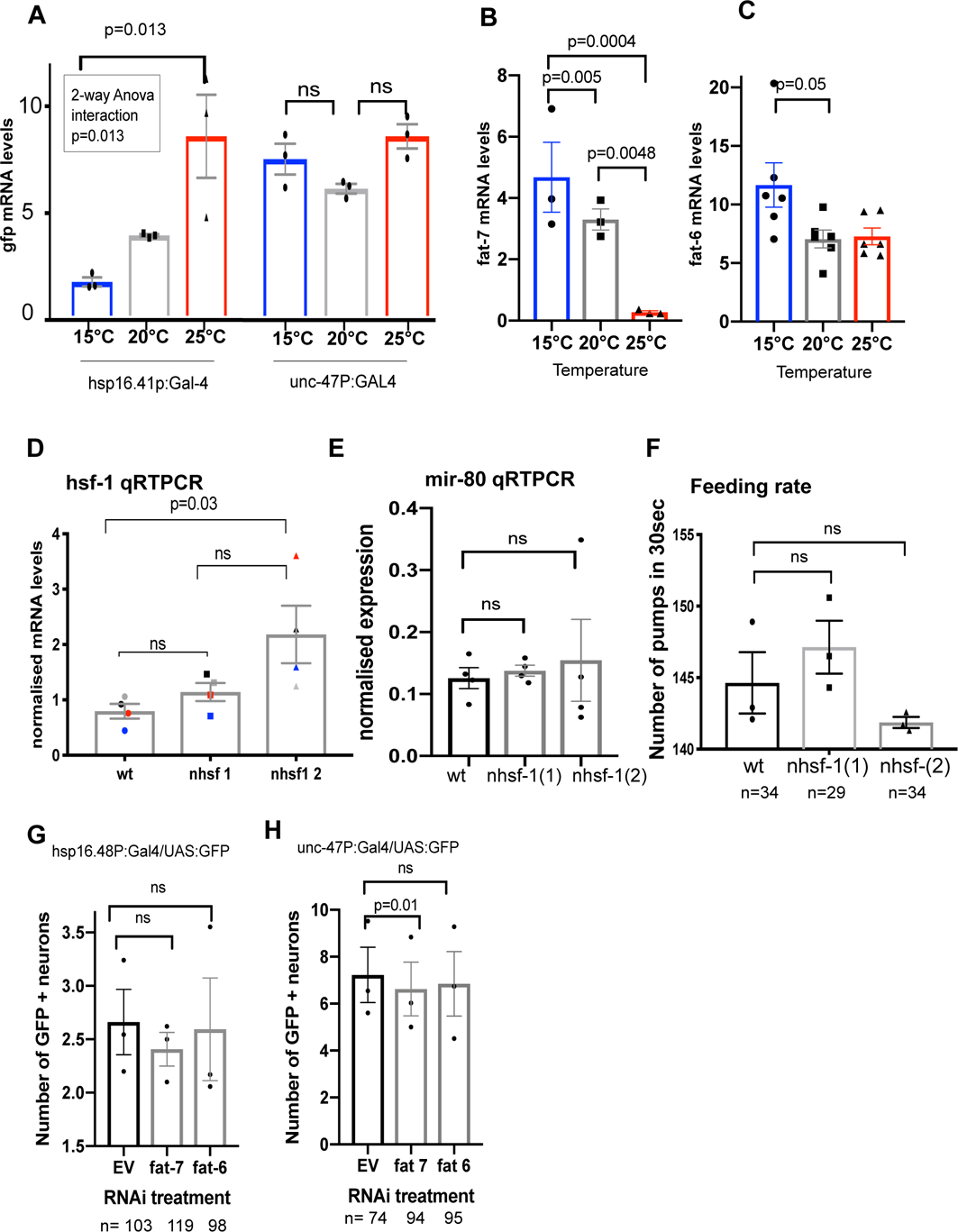
(**A**) Two-way ANOVA shows a significant difference in *gfp* mRNA levels in animals raised at 16℃ vs 25℃ when *gfp* is driven by *hsp-16.41* promoter and not when it is driven by the *unc-47* promoter. A 2-way ANOVA shows significant (p=0.022) interaction between temperature and genotype, indicating that the difference in *gfp* levels between 16℃ and 25℃ is higher for the promoter *hsp16.41* than for *unc-47* (Sup Table S1A). The presence of the GAL4/UAS system *per se* does not cause a temperature dependent change in *hsp16.41* transcriptional output. (**B-C**) Bulk qPCR, showing that the expression of fat desaturases *fat-7* (**D**) and *fat-6* (**E**) inversely correlates with temperature. *fat-6* expression levels significantly decrease by 39% from 16℃ to 20℃ but undergo no further change from 20℃ to 25℃. *fat-7* expression levels decrease by 46% decrease from 16℃ to 20℃, and by 93% decrease from 20℃ to 25℃. Each dot represents a biological replicate and the bars represents the standard error. Statistics were performed using one-way ANOVA (Sup Table S1B). **D)** *hsf-1* mRNA expression levels, detected by qRTPCR in wild type and *hsf-1* strains that over-express *hsf-1* in neurons. Paired data has been colour-coded to highlight the consistent trend of *hsf-1* up-regulation in *nhsf-1*(1), although this is not significant. *nhsf-1*(2) has significantly increased levels of *hsf-1.* Statistics was performed using a One-way ANOVA test (Sup Table S1C). (**E-F**) Ectopic expression of *hsf-1* in neurons does not cause starvation. (**E**) qRTPCR of a known starvation marker, *mir-80*, which is not upregulated in nHSF-1 animals. (**F**) animals have normal levels of pharyngeal pumping, which indicates they are feeding normally. n values are indicated at the bottom of the figure. Each dot represents a biological replicate, statistics was performed using One-way ANOVA (Sup Table S1D). (**G-H**) Reduced *fat-6* or *fat-7* levels do not induce neuronal stress, indicating that there is no double feedback loop. Animals were treated with empty vector (EV), or with *fat-7* or *fat-6* RNAi. The number of GFP positive neurons was counted for animals bearing (**G**) *hsp16.48p*:Gal4;UAS:GFP; and (**H**) *unc-47*p:Gal4; UAS:GFP. Each dot represents the results of a biological replicate and statistics was performed using One-way ANOVA. The number of animals analysed across replicates (n) is expressed at the bottom of the figure (Sup Table S1E). Bars represent standard error of the mean. All data and statistics have been summarised in Sup Table S1A-D.

**Figure S2.**
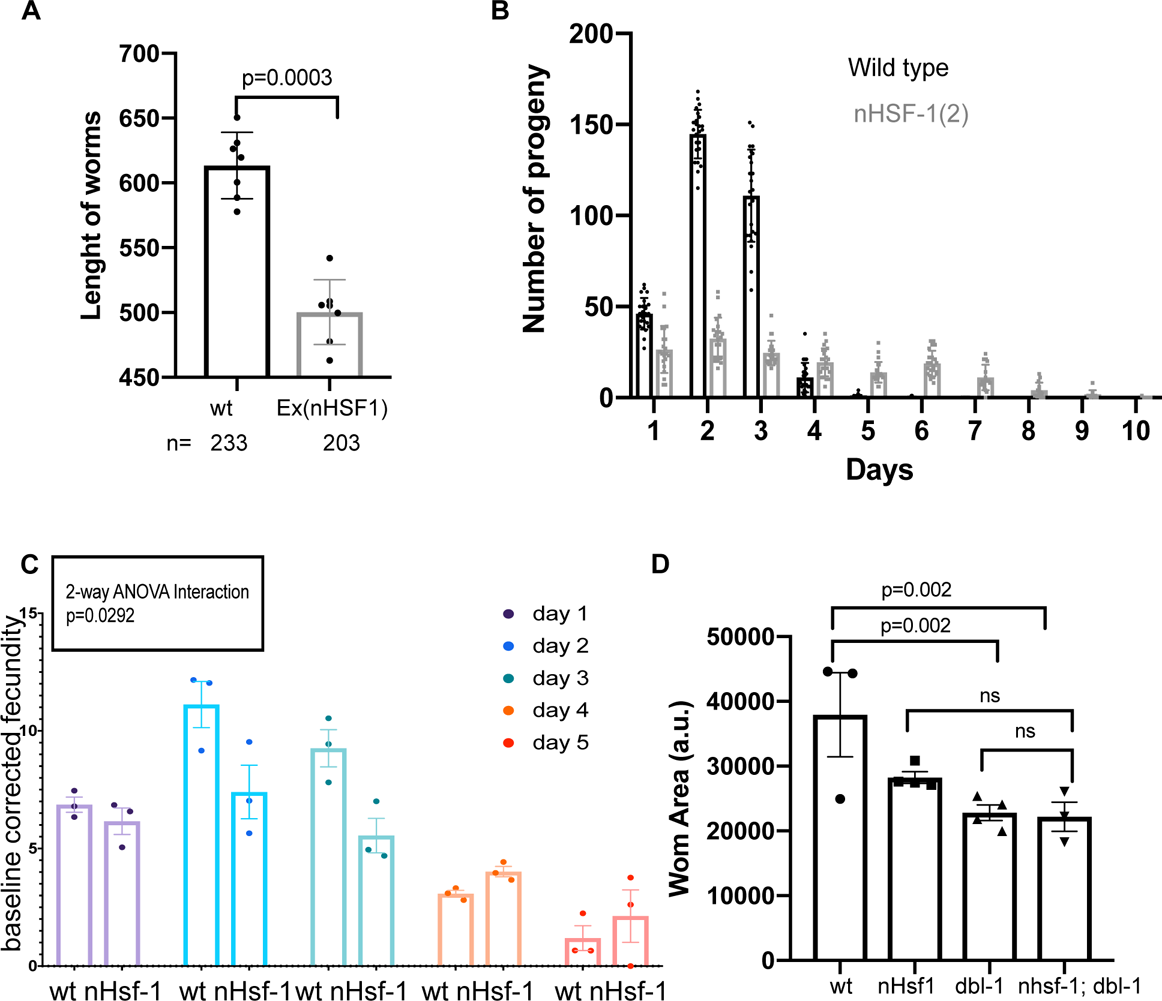
nHSF-1 phenocopies BMP mutants. **(A)** Length of the major axis of young adult *C. elegans* of the genotype *uthEx663[rab-3p::hsf-1; myo-2p::tomato] Is[fat-7p::GFP]*.). Siblings that contain the over-expression construct were distinguishable by a red signal in the pharynx (produced by the *myo-2p::tdtomato* co-injection marker). The length of the major axis in wild type animals (which do not contain the extra-chromosomal array) is on average 0.6 mm. In nHSF1 animals (which contain the extrachromosomal array), this length is significantly reduced by 20%, relative to WT. Each dot represents a biological replicate and n values are indicated at the bottom of the figure. The bar is the SE of the mean, and statistics was performed using the paired t-test (Sup Table S2A). Note that the Y axis does not start from 0. (**B-C**) Ectopic nHSF1 expression causes a germline senescence phenotype. (**B**) Number of self-progeny laid per day at 20℃, from day 1 to 10 of adulthood, quantified in one experiment. Each dot represents the total progeny output of a single worm. nHSF1(2) animals produce fewer progeny than do wildtype worms in the first three days, but continue laying eggs after day 5, when WT animals cease to reproduce. (**C**) Fertility of self-reproducing animals for 3 biological replicates of wild type and nHSF1(2). Each dot represents the average progeny of multiple animals per replicate. The statistical test was corrected for the heterogeneity of variance using sqrt transformation. A 2-way ANOVA test shows a positive interaction, indicating that the difference in reproductive output varies significantly: it is lower than WT during the first 3 days of adulthood and then remains higher than wild type for the last two tested days (Sup Table S2B). (**D**) The small size of nHSF-1 is caused by reduced BMP signalling. Area in arbitrary units (a.u.) of the genotypes. BMP is epistatic to nHSF-1 because the double mutants are not shorter than any single mutant. nHsf-1 causes a 26% reduction in size, *dbl-1(nk3)* causes a 39% reduction in size, and the double mutant is 41% smaller than wild type. The double mutants are not significantly different in size from any single mutant. Each dot corresponds to a biological replicate, n values were all larger than 62 per genotype. Error bar corresponds to the standard error of the mean, and statistics were performed using a one-way ANOVA test. All data and statistics have been summarised in Sup Table S2.

**Figure S3.**
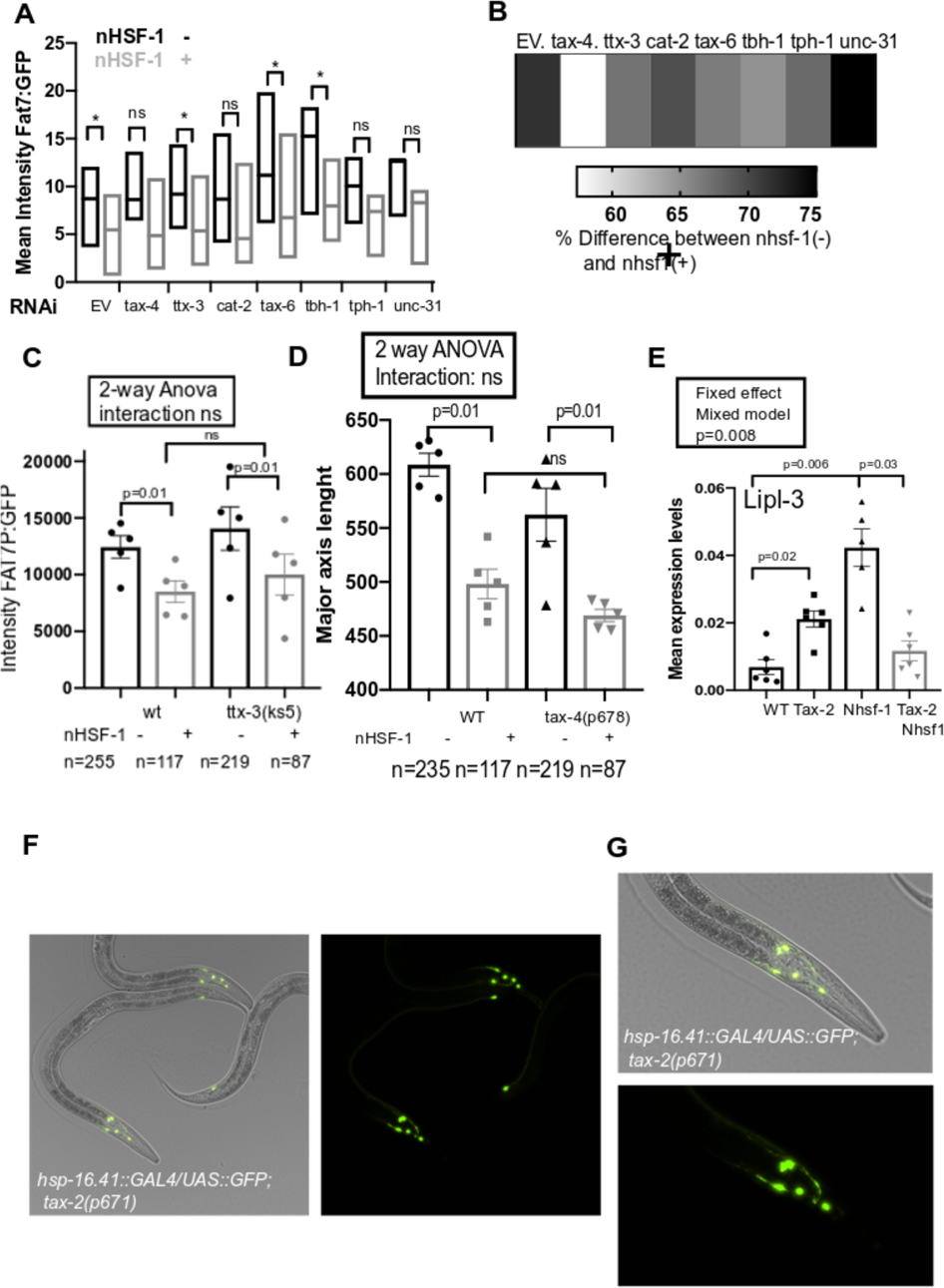
Neuronal modifiers of fat metabolism in nHSF-1. **(A)** A RNAi-based screen for suppressors of the repressive effect of neural-Hsf-1 on *fat-7*p: GFP expression. The screen was performed using MOC194 (*rab-3*P:*hsf-1* (*uthEx663[rab-3p::hsf-1; myo-2p::tdtomato]; nIs590 [fat-7p::fat-7::GFP + lin15(+)] V*). The graph shows the comparison between the effect of an empty vector (EV) with the knock down of 8 genes expressed in neuronal cells that are known or suspected to influence fat metabolism. The Y axis shows the levels of GFP measured by fluorescence microscopy and driven by the *fat-7* promoter (+/-SEM). Siblings containing the over-expression construct were distinguished by the presence of a red signal in the pharynx (coming from *myo-2p::tdtomato* co-injection marker). The Y axis represents the average mean intensity of GFP from 3 replicates. The KOs that result in a significant suppression of *fat-7*p:GFP repression by nHSF1 are *tax-2*, *cat-2*, *tax-6*, *tbh-1* and *unc-31*. However, all of these also cause an increase in *fat-7*pGFP in controls animals, with the exception of *tax-6* and *tax-4* (Sup Table S3A). **(B)** Percentage change among pairs treated with RNAi treatment shows that *tax-4* causes the largest rescue of fat-7p:GFP, where for the control pair, the difference between EV and nhsf-1 is 37%, this difference is only 19% when animals are fed *tax-4* RNAi. **(C)** Mean Intensity of GFP driven by the *Fat-7* promoter is not changed in the absence of a functional thermosensory neurons. The graphs show Mean Intensity GFP levels for control animals compared to both *ttx-3 (ks5)* (**C**). Two-way ANOVA shows a negative interaction, indicating that mutations in *ttx-3* or *gcy-8* do not change GFP levels in the presence of nHSF1 (Sup Table S3B). **(D)** 2-Way ANOVA interaction indicates that the smaller size of nHSF-1 animals is not rescued by *tax-4(p678)* (Sup Table S3C). **(E)** Relative expression levels of *lipl-3* mRNA by qRTPCR in wild type, *tax-2(p671),* nHSF-1(2) and *tax-2(p671);* nHSF1(2) double mutant animals. The levels of *lpl-3* are suppressed in the absence of *tax-2* function (Sup Table S3D). (**F-G)** *tax-2(p671)* mutants are able to express *hsp-16.41* in neurons. Young adult *hsp16.41p*:Gal4/UAS:GFP;tax-2(p671) animals raised at 20℃. 20X overlay of GFP fluorescence and Nomarski images of a GFP is expressed exclusively in anterior neuronal cells with axons.

**Figure S4.**
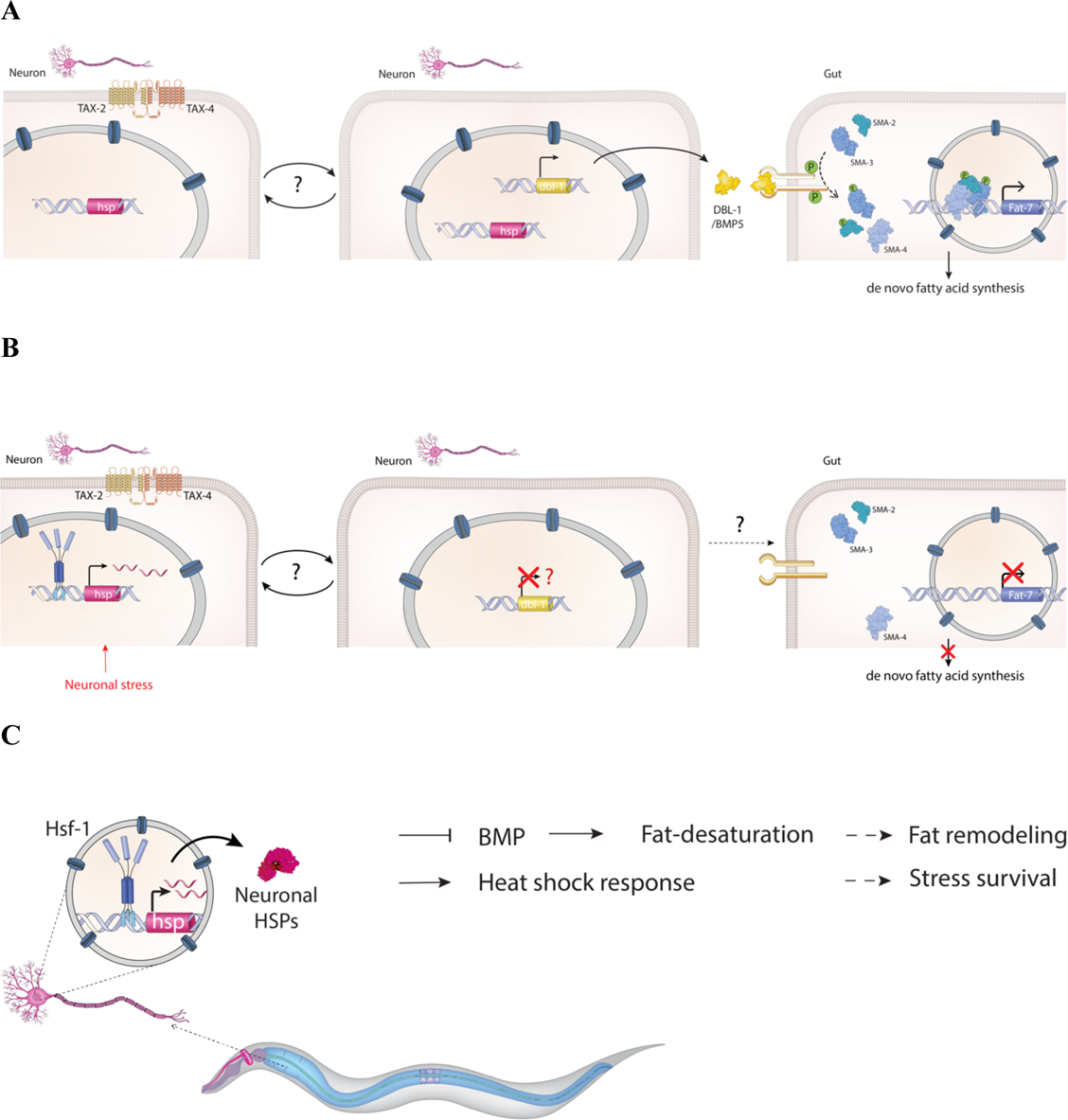
Model of how neuronal stress modulates fat metabolism in distant fat storage cells. (A) In wild type animals, TGFβ ligand (DBL-1/BMP5) is secreted from neurons and activates the Sma/Mab BMP signalling pathways by phosphorylating SMADs proteins. SMADs transcriptionally regulate the expression of genes involved in LD in fat storage cells (gut and hypodermis), including Fat-7. The binding of SMADs directly to Fat-7 promoters is hypothetical and has not been directly tested. (B) The activation of the HSR in neurons expressing the cGMP receptor TAX-2/TAX-4 directly or indirectly dampens BMP signalling with consequences for the expression of *fat-7* and other genes involved in metabolism in fat storage cells. (C) Ectopic neuronal stress de-activates fat desaturation causing profound fat remodelling changes in peripheral tissues.

**Figure S5.**
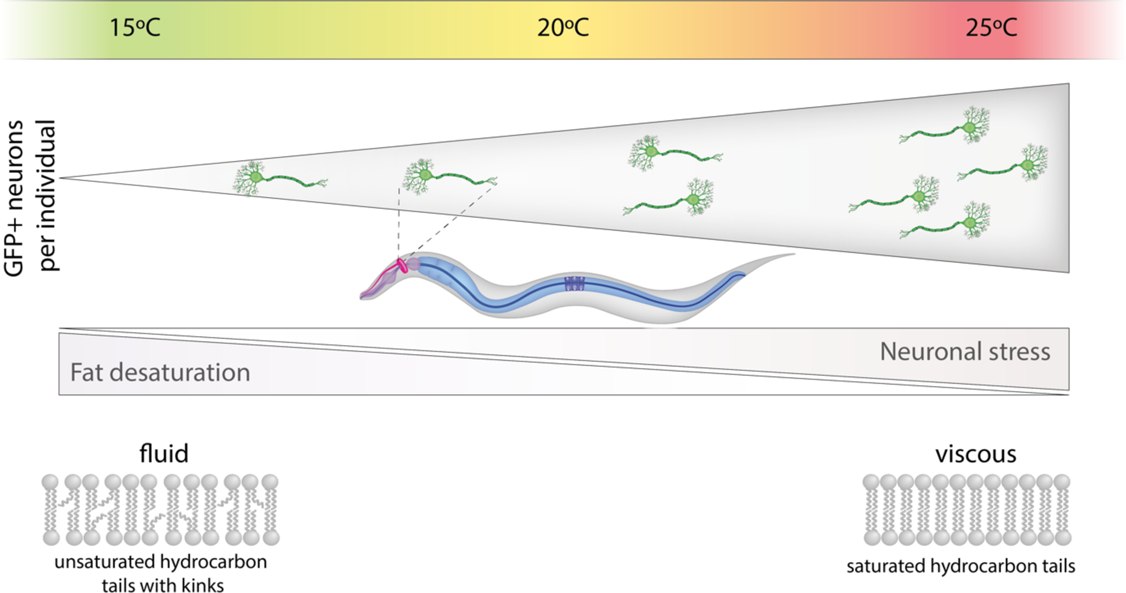
Thermostat senses ambient temperature and modulates membrane fluidity. Ectotherms rely on homeoviscous adaptation (HVA) to survive temperature fluctuations. Consistent with HVA, the FA composition of membranes can be modulated by fat desaturase enzymes. Modulating the number of double bonds in the hydrocarbon tails of FAs, tunes membrane fluidity to ambient temperature, providing appropriate thermodynamic conditions for biochemical reactions to occur. HSF-1 responds to warm temperature by activating a stress response in sensory neurons. When the stress response is very high or made constitutive (by ectopic expression of Hsf-1) the thermostat switches fat metabolism to make membranes viscous and better adjusted to warm temperatures.

